# Escherichia coli grown in cost-effective conical tubes produces more plasmid DNA

**DOI:** 10.1101/2024.03.13.584835

**Authors:** Mayra Marquez, Qian Chen, Silvia Cachaco, Hongyan Sui, Tomozumi Imamichi

**Affiliations:** Laboratory of Human Retrovirology and Immunoinformatics, Frederick National Laboratory for Cancer Research, Frederick, Maryland, 21702, USA

## Abstract

Before the COVID-19 pandemic, we grew plasmid-transformed *Escherichia coli* in Falcon round-bottom-polypropylene tubes (F-Round-PP) and isolated plasmid using a Miniprep kit. When obtaining sufficient quantities of F-Round-PP became problematic during the pandemic and the inflation, we grew them using any available tubes. Notably, we observed that plasmid yield from cells grown in a cost-effective Oxford conical-polypropylene tubes (O-Conical-PP) was higher than that from F-Round-PP. We assessed the impact using O-Conical-PP and other conical brand tubes. As a result, the plasmid yield from O-Conical-PP (the list price is nearly 1/3 of F-Round-PP) was 1.5-fold higher than other PP tubes (p<0.001). We propose that researchers may need to re-assess the effectiveness of their laboratory supplies to optimize the budget during this inflationary period.

## Introduction

During the COVID-19 pandemic, research progress in basic science laboratories, including those involved in unrelated COVID-19 studies, was delayed. One of the reasons for this was the delay in receiving lab supplies owing to a shortage of materials and an increase in prices, which affected research budgets.

Our lab has conducted molecular cloning of Human Immunodeficiency Virus type-1 (HIV-1) from blood plasma of people living with HIV infection using standard molecular biology methods (e.g. reverse transcription, polymerase chain reaction (PCR), transformation and Sanger sequencing) [1, 2]. Before the COVID-19 pandemic, we routinely grew *Escherichia coli* (*E. coli*) containing plasmid DNA in Luria-Bertani (LB) media using round bottom polypropylene (PP) tubes, and extracted the plasmid DNA using a MiniPrep Kit [3]. These tubes were optimal because PP has a high tensile strength and impact resistance and exhibits good resistance to fatigue and stress cracking [4, 5]. In contrast, polystyrene (PS) is relatively brittle, has lower impact resistance than PP, and is more prone to stress cracking under high-speed centrifugation [6, 7]. When obtaining sufficient quantities of Round-PP became difficulty during the COVID-19 pandemic and the inflation, we grew the cells using any available tubes, regardless of manufacturers and material. Notably, we observed that the plasmid DNA yield from the cells grown in low-cost conical PP tubes was higher than that of the cells grown in Round-PP. In this report, we evaluated the impact of tubes materials and shape on plasmid DNA yield.

## Materials and Methods

### Isolation of plasmid DNA

Glycerol stock of *Escherichia coli* (*E. coli*) (TOP10 strain, Invitrogen, Thermo Fisher Science, Waltham, MA, USA) transformed with plasmid pCR2.1 (Invitrogen) containing 3.6 kbp DNA fragment (a total DNA size is 7.5 kbp) [8], or Stbl3 (Invitrogen) with plasmid pUC19 (Invitrogen) containing 12.8 kbp DNA (total size is 15.5 kbp) [9] was streaked on Luria-Bertani (LB) media (Quality Biological, Germantown, MD, USA)-Bacto Agar (Becton Dickinson, Franklin Lakes, NJ, USA) plate containing 100µg/ml ampicillin (MilliporeSigma, Burlington, MA, USA) (LA-plate), Five single colonies were individually inoculated in 5 mL of LA medium in 50 mL Falcon polypropylene (PP) tube (Cat# 352098) and then cultured for overnight at 37°C at 225 rpm. The next day, one % of the overnight culture was reinoculated into tubes containing warm LA (1/5 vol of each tube capacity; detail of the volume is described in the results and discussion) and cultured for 16 hr at 37°C at 225 rpm, and then, 1 mL of each bacteria suspension was subjected to isolate plasmid DNA using the S.N.A.P. Plasmid DNA MiniPrep Kit (Invitrogen). *E. coli* was pelleted in a 1.5 mL microcentrifuge tube (Eppendorf, Cat# 022363654) at 8000 rpm for 1 min at room temperature (RT), and then LB media was removed by decant. The cell pellets in each tube were resuspended in 150 µL of the Suspension buffer (Invitrogen) by pipetting up and down, and then 150 µL of the Lysis butter was mixed and incubated for 3 min at RT. After the incubation, 150 µL of the Precipitation buffer (Invitrogen) was added to each tube and mixed six-time by slow inversion. After confirming the formation of white aggregation, the tubes were centrifuged at 15,000 rpm for 5 min at RT. After centrifugation, supernatant (approximately 450 µL) was mixed with 600 µL of the binding buffer (Invitrogen) by pipetting up and down, and the loaded onto a spin column (Invitrogen) on a collection tube (Invitrogen). The columns were centrifuged at 6000 rpm for 1 min, and the flow-through was discarded. The columns were washed with 500 µL of the Wash buffer (Invitrogen) at 6000 rpm for 1 min on the collection tube and then further washed with 900 µL of the Final wash buffer (Invitrogen) for 1 min at 6000 rpm. The columns were put on new 2 mL collection tubes (Qiagen) and centrifuged at 15000 rpm for 1 min to dry the columns. The dried columns were set onto 1.5mL new microcentrifuge tubes, and 50 µL of molecular grade water (Ambion, Thermo fisher Science) was added and incubated for 3 min at RT, and then plasmid was eluted from the columns by centrifugation at max speed for 1 min.

### Quantification of DNA and bacterial growth

Bacterial growth was measured by optical density (OD) at A600 nm using SpectraMax M5 (Molecular Devices, San Jose, CA, USA). Briefly, 200 µL of suspension was put into a 96-well plate (Corning, Cat# 3598) in duplicate and then OD was measured. The average of OD for each bacterial suspension was used as bacterial growth values. DNA yield was determined by quantifying A280 and A260 using a Nanodrop (Thermo Fisher Scientific) and obtained µg/mL of DNA concentration and the A260/A280 ratio. The DNA concentration was divided by each A600 to compare DNA yield among samples, and the results were demonstrated as a DNA/A600 ratio.

### Statistical analysis

To demonstrate statistical significance, one way ANOVA (Prism version 10.2.0, GraphPad, La Jolla, CA, USA) was used, and each figure were created using the PRISM.

## Results and Discussion

To evaluate whether materials or tube shapes significantly affect plasmid DNA yield and potentially save lab budget, we conducted five independent isolations from five independently inoculated cell growth by five researchers using five different tubes: Falcon round polypropylene tube (F-Round-PP, Cat# 352059), Oxford conical PP tube (O-Conical PP, Cat #OCT-15B), Fisher Brand conical PP tube (FB-Conical-PP, Cat#.05-539-5), Corning conical PP tube (C-Conical-PP, Cat # 430052), and Falcon conical polystyrene tubes (F-Conical-PS, Cat# 352095) using the MiniPrep Kit as described in the Materials and Methods. As the cap sealing system of each tube is unique, e.g., dual position snap-cap plug seal or plug seal caps, thus it was presumed that the airflow might affect the difference in DNA yield; therefore, during incubation, the cap was set on the tube without closing and taped on it to avoid evaporation. In addition, to retain air permeability and heat conductivity in each tube in the same condition, LB media was added at 1/5 of the tube capacity (2. 8 mL for F-Round-PP, and 3 mL for other tubes). The *E. coli* growth was monitored for optimal densities at 600 nm (A600). As a result, the cell growth was not significantly different among all tested samples containing 7.5kbp plasmid DNA (Figure 1A). 1 ml of cell suspension from each tube was used to isolate plasmid DNA as described in the Materials and Methods, and the quantified DNA amount was normalized to A600. Although A260/A280 ratio was similar among all samples (1.89 ∼1.90), of interest, we found that the amount of plasmid DNA/A600 obtained from O-Conical-PP tubes was nearly1.5-fold higher than that from F-Round-PP (p <0.01, Figure 1B) and FB-Conical-PP or C-Conical-PP (p <0.001). Notably, DNA yield (DNA/A600) from F-Conical-PS was similar to that from O-Conical-PP.

**Figure 1.**
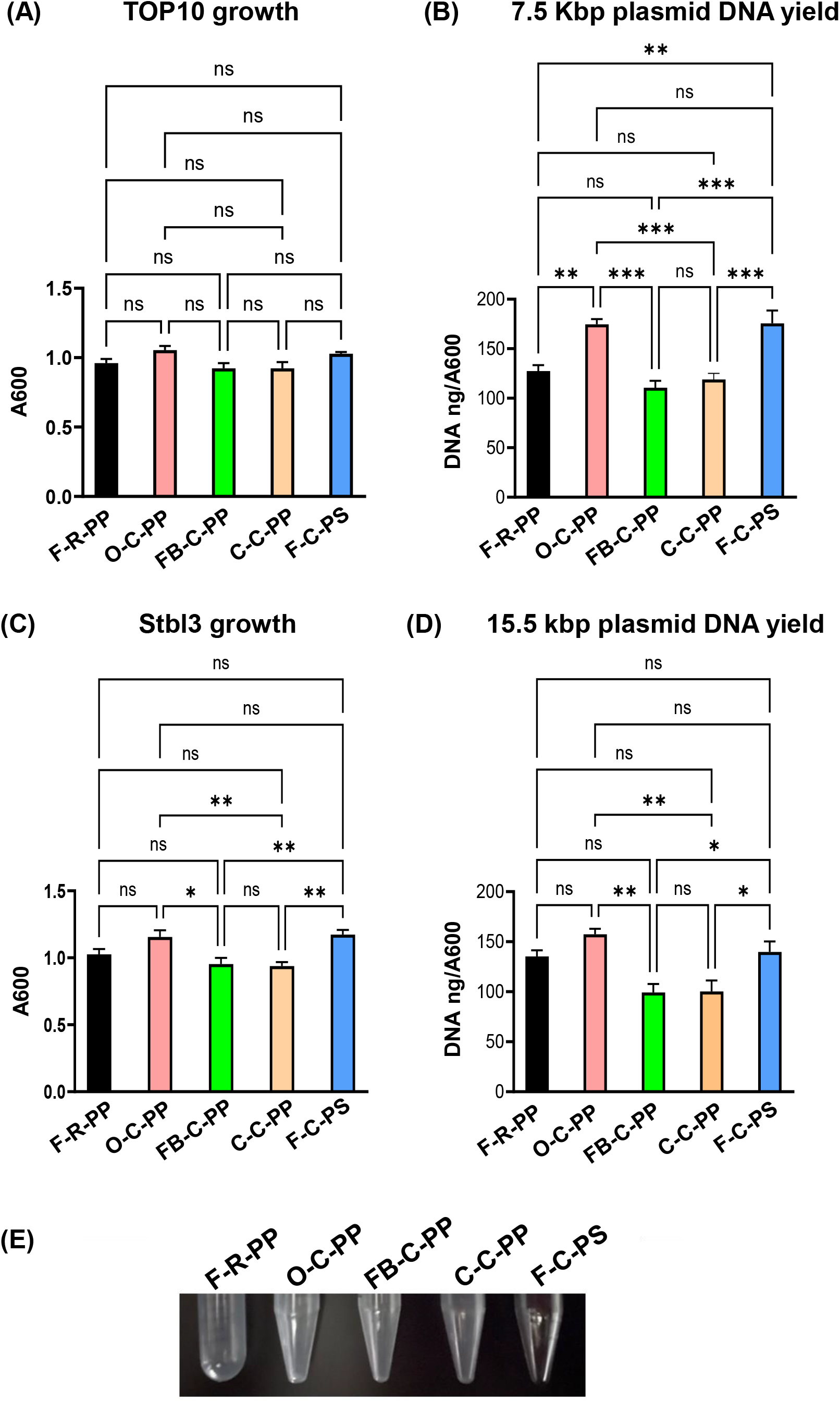
Comparison of *E. coli* growth and plasmid DNA yields among different tubes. TOP10 containing 7.5 kbp plasmid DNA (**A** and **B**) or Stabl3 containing 15.5 kbp plasmid DNA (**C** and **D**) were cultured in 2.8 mL or 3 mL (1/5 of each tube capacity) of LB-medium supplemented with 100 µg/mL ampicillin (Sigma Aldrich, St. Louis, MO, USA) in five tubes (F-R-PP: F-Round-PP, O-C-PP: O-Conical-PP, FB-C-PP: FB-Conical-PP, C-C-PP: C-Conical-PP, F-C-PS: F-Conical-PS) for 17 hr at 37°C with 2200 rpm. (**A** and **C**) Cell growth was quantified by measurement of the optical density at 600 nm. (**B** and **D**) Plasmid DNA was isolated from one mL of suspension from each tube using the S.N.A.P. Plasmid DNA MiniPrep Kit and eluted in 50 uL of molecular biology grade H_2_O (Ambion, Thermo Fisher Science). DNA was quantified using Nanodrop (Thermo Fisher). The DNA amounts were normalized to A600 nm. Total five independent assays were run using five independent inoculated cell suspension. The results indicate mean and ±SE. A statistical analysis was conducted with one way ANOVA using Prism version 10.2.0 (GraphPad, Boston, MA, USA). n.s.: not significant, *: *p*<0.05, **: *p*<0.01, ***: *p*<0.001. **(E)** Comparison of the bottom shape of each tube. An image of the bottom shapes of the five tubes (F-R-PP: F-Round-PP, O-C-PP: O-Conical-PP, FB-C-PP: FB-Conical-PP, C-C-PP: C-Conical-PP, F-C-PS: F-Conical-PS) was taken.

To determine whether the increased output effect was *E. coli* strain- or plasmid size-dependent, we compared extraction of a 15.5 kbp plasmid from the STABL3 strain (Invitrogen) [9]. STABLE3 growth was similar levels in F-Round-PP, O-Conical-PP, and F-Conical-PS and the amounts of plasmid DNA from those three tubes were comparable (Figure 1C). Cells grew less in FB-Conical-PP and C-Conical-PP tubes than in O-Conical-PP tubes. Subsequently, the amounts of plasmid DNA from those tubes were less than in O-Conical-PP (Figure 1D). These data indicated that we tend to obtain more DNA from *E. coli* grown in O-Conical-PP tubes than others, regardless of strain and plasmid size.

At this moment, we do not know the difference in the production process of each PP tube and the precise tubing specifications (inner diameter and the wall thickness on each tubing) for each tube that may affect the surface area and thermal conductivity. However, as shown in Figure 1E, the bottom shape of O-Conical-PP tube is flat-tipped, while others have pointed ends, and O-Conical-PP tube is less transparent than other PP tubes. This difference might influence thermal conductivity, oxygen diffusion, liquid convection, which resulted in increased plasmid DNA without affecting cell number in growth. As of March 2024, the list price is US $135 for 500 O-Conical-PP tubes, US $425 for 500 F-Round-PP, US $353 for FB-Conical-PP, $480 for C-Conical-PP tubes, or US $414 for F-Conical-PS. Using O-Conical-PP tubes, we required less LB media to obtain more DNA with a smaller budget than other PP tubes. We believe that this finding regarding the difference in plasmid yield significantly impacts savings in laboratory budgets and LB. During the current inflationary period, we encourage researchers to re-assess the cost and effectiveness of their laboratory supplies to optimize budget use.

## Financial disclosure/Acknowledgements

This project was funded in whole or in part with federal funds from the National Cancer Institute, National Institutes of Health, under Contract No. HHSN261200800001E. This research was supported (in part) by the National Institute of Allergy and Infectious Diseases. The authors thank HC. Lane, HA. Young and Y. Sei for discussing the project, and W. Chang and S. Laverdure for critical reading.

## Ethical conduct of research

N/A (this paper contains none of data involving humans or animals)

